# Machine Learning Identification of Functional Trait Syndromes Associated with Responsiveness to Arbuscular Mycorrhizal Fungi

**DOI:** 10.64898/2026.10.05.756734

**Authors:** Tariq Shah, Mohamed Akram Errahmouni, Mohamed Hijri

## Abstract

Arbuscular mycorrhizal fungi (AMF) are widespread plant symbionts that enhance nutrient acquisition, growth, and stress tolerance, yet plant responsiveness to AMF varies substantially and remains difficult to predict. We developed Trait2Myco, a Random Forest based machine-learning framework that integrates plant functional traits and environmental variables to predict mycorrhizal growth response (MGR). The framework was trained using 80 observations spanning four grass species grown under four precipitation regimes. Predictor variables included leaf and root economic traits together with precipitation, while total MGR served as the response variable. A full model incorporating ten functional traits and precipitation was trained using 80% of observations and evaluated on an independent testing dataset (20%, n = 16). The full model explained 41% of variation in MGR (R² = 0.41), whereas a reduced ecological model containing only specific leaf area, specific root area, root carbon concentration, and precipitation improved predictive performance and explained 51.7% of variation (R² = 0.517). Ten-fold cross-validation of the full model yielded an R² of 0.983, RMSE of 0.046, and MAE of 0.036. Variable importance analysis identified specific leaf area, precipitation, root carbon concentration, and specific root area as the strongest predictors of responsiveness. Partial dependence analysis revealed nonlinear relationships between these predictors and MGR, while principal component analysis showed that highly responsive plants occupied distinct regions of functional trait space characterized by resource-acquisitive trait syndromes. Leaf and root traits contributed nearly equally to model performance, accounting for 41.9% and 41.3% of cumulative importance, respectively, while environmental factors contributed 16.8%. An exploratory analysis using an independent AMF colonization dataset produced similarly strong predictive relationships, suggesting that trait-based machine-learning approaches may generalize across multiple AMF-associated responses. Trait2Myco provides a reproducible framework for identifying plant trait syndromes associated with responsiveness to AMF and offers a practical tool for predicting mycorrhizal benefits across environmental gradients.

**Graphical Abstract:** 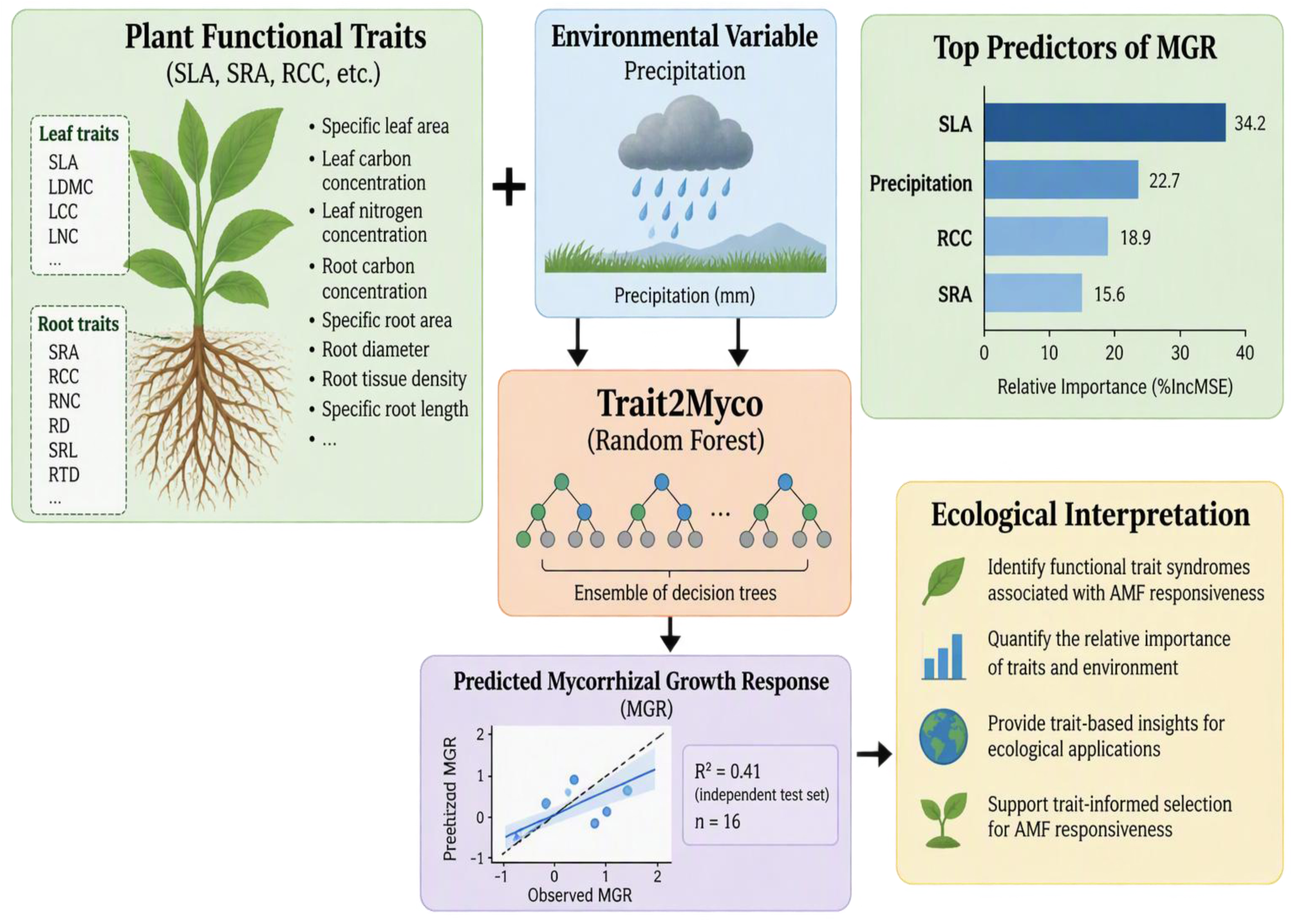

## 1. Introduction

Arbuscular mycorrhizal fungi (AMF) form one of the most ancient and widespread mutualistic associations in terrestrial ecosystems, colonizing the roots of many vascular plant species and playing a central role in global nutrient cycling (Smith and Read 2008; Brundrett 2009; Bonfante and Genre 2010; van der Heijden et al. 2015). Through this symbiosis, AMF enhances phosphorus and nitrogen acquisition, improves water relations, increases tolerance to abiotic and biotic stress, and contribute substantially to ecosystem productivity (Smith and Read 2008; van der Heijden et al. 2015; Begum et al. 2019). In exchange, host plants allocate a significant fraction of recently fixed carbon to their fungal partners, establishing a reciprocal exchange that structures both plant performance and belowground community function (Smith and Read 2008; Bever et al. 2009; van der Heijden et al. 2015).

Despite the near-universal distribution of AM symbiosis, the magnitude and even the direction of plant growth response to AMF colonization vary enormously among species, genotypes, developmental stages, and environmental conditions (Janos 1980; Johnson et al. 1997; Klironomos 2003). This variation is commonly quantified as the mycorrhizal growth response (MGR), the proportional difference in plant performance between inoculated and non-inoculated (or weakly colonized) plants. Responses span a continuum from strongly positive to neutral to clearly negative (Johnson et al. 1997; Klironomos 2003), and this variability has made MGR difficult to predict from colonization intensity alone, since the two are frequently decoupled (Smith and Smith 2011; Hart and Reader 2002). Identifying the drivers of this variation is a central problem in mycorrhizal ecology, with direct consequences for whether inoculation-based strategies succeed in agriculture and ecological restoration (Hoeksema et al. 2010; Treseder 2013; Lehmann et al. 2012).

A range of mechanisms have been proposed to explain variation in MGR, including nutrient supply demand dynamics (Koide 1991), carbon-cost efficiency of the symbiosis (Corrêa et al. 2011), host phylogeny and fungal community identity (Maherali and Klironomos 2007; Klironomos 2003), soil nutrient availability and abiotic context (Hoeksema et al. 2010; Treseder 2013; Veresoglou et al. 2012), and root system architecture (Wilson and Hartnett 1998; Hart and Reader 2002). More recently, plant functional traits have emerged as a promising integrative framework for explaining this variability. The leaf economics spectrum describes a continuum from resource-acquisitive species, characterized by high specific leaf area and rapid nutrient turnover, to resource-conservative species with denser, longer-lived tissue (Wright et al. 2004; Reich 2014). An analogous economic spectrum exists belowground, where root traits such as specific root length, root diameter, and tissue density reflect trade-offs between independent nutrient foraging and reliance on fungal symbionts (Roumet et al. 2016; Bergmann et al. 2020). Notably, root economic variation does not collapse onto a single acquisitive–conservative axis: Bergmann et al. (2020) demonstrated that a distinct collaboration gradient, orthogonal to the conservation gradient, captures the degree to which species invest in traits favoring mycorrhizal partnership over independent root foraging. Building on this framework, Chagnon et al. (2013) proposed that both plant and fungal traits can be organized along life-history axes that jointly predict the ecological outcome of the symbiosis, and subsequent work has reinforced the view that leaf and root economic traits jointly shape both the degree of AMF colonization and the benefit a host ultimately derives from it (Freschet et al. 2021; Delavaux et al. 2017).

Despite this conceptual progress, predicting MGR from trait data in practice remains difficult, because trait effects on mycorrhizal responsiveness are frequently nonlinear, context-dependent on the abiotic environment, and the product of interactions among multiple traits rather than any single trait acting in isolation (Chagnon et al. 2013; Hoeksema et al. 2010). Conventional linear statistical models are poorly suited to capturing this complexity and typically require a priori specification of functional form and interaction terms that are rarely known beforehand in ecological trait datasets. Machine-learning algorithms offer a promising alternative: Random Forest models in particular can accommodate nonlinear relationships, high-order interactions among predictors, and the multicollinearity that is endemic to correlated trait datasets, all while generating quantitative, interpretable measures of variable importance (Breiman 2001; Cutler et al. 2007; Prasad et al. 2006). Machine-learning approaches have seen growing use across plant and soil ecology (Cutler et al. 2007; Prasad et al. 2006), yet predictive frameworks specifically designed to identify the functional trait syndromes associated with mycorrhizal responsiveness remain limited.

To address this gap, we developed Trait2Myco, a Random Forest–based machine-learning framework that integrates leaf and root functional traits with an environmental covariate to predict plant responsiveness to arbuscular mycorrhizal fungi. Using 80 observations spanning four plant species and four precipitation regimes, we examined whether leaf and root economic traits could predict variation in MGR, asking specifically: (i) which functional traits contribute most strongly to responsiveness; (ii) whether a reduced set of ecologically meaningful traits can explain a substantial proportion of variation in MGR; and (iii) how highly responsive and weakly responsive plants are distributed within multivariate trait space. To evaluate whether trait-based prediction generalizes beyond the traits used in model training, we additionally applied the framework to an independent dataset relating host growth traits to AMF root colonization intensity (Zhao et al. 2026; Werner and Kiers 2015). By combining machine learning with ecological interpretation across two independent datasets, Trait2Myco provides a reproducible framework for understanding trait-mediated variation in plant responses to AMF and for identifying species likely to benefit from mycorrhizal symbiosis.

## 2. Materials and Methods

### 2.1. Development of the Trait2Myco framework

Trait2Myco was developed as a Random Forest based machine-learning framework to identify plant functional trait syndromes associated with responsiveness to AMF. The framework integrates ecological trait data and an environmental covariate to predict MGR and to identify the most influential drivers of responsiveness. Random Forest was selected because it models nonlinear relationships, accommodates interactions among predictors without pre-specification, and remains robust to the multicollinearity commonly encountered among correlated functional traits (Breiman 2001; Cutler et al. 2007). The overall workflow of Trait2Myco, including data integration, model development, prediction, and ecological interpretation, is illustrated in Figure 1.

**Figure 1.**
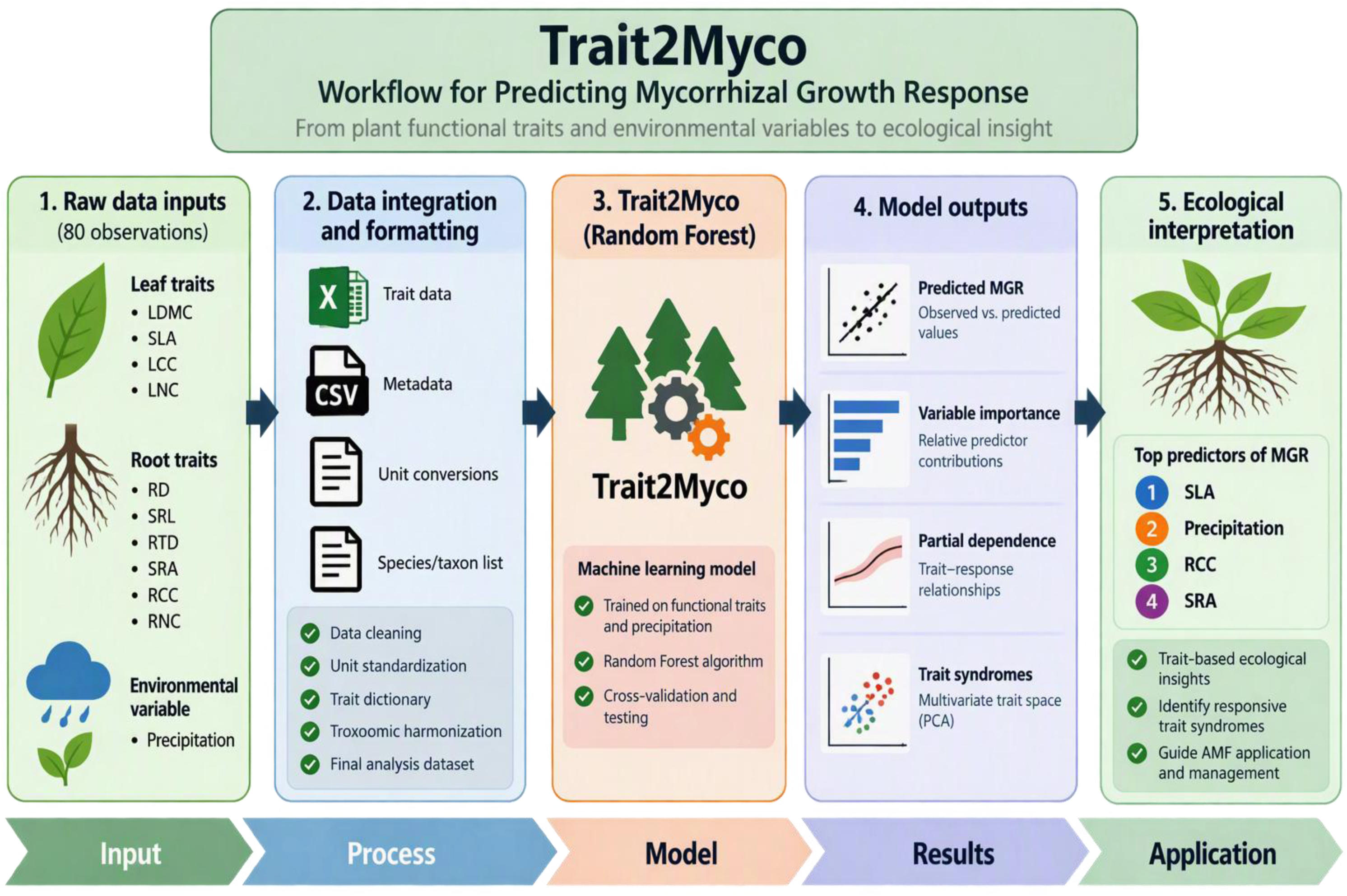
Conceptual overview of the Trait2Myco framework for predicting mycorrhizal growth response from plant functional traits and environmental variables.

### 2.2. Primary dataset: sources and preparation

The primary dataset was obtained from the Dryad Digital Repository (Zhao et al. 2026; https://doi.org/10.5061/dryad.3j9kd51v3), comprising measurements from four plant species exposed to four precipitation regimes, with five biological replicates per species–precipitation combination (80 observations total after merging). Component data files like plant biomass and shoot nutrients, mycorrhizal response measurements, and leaf and root functional traits were merged using species identity and replicate number as shared identifiers, producing a single dataset containing both trait values and response measurements for every observation. Functional trait and environmental data were integrated and used as inputs into the Trait2Myco Random Forest framework, which generated predictions of mycorrhizal growth response (MGR) and associated model outputs, including variable importance rankings, partial dependence analyses, and trait-space ordinations (Fig. 1).

### 2.3. Functional trait predictors

Trait2Myco incorporated leaf and root traits representing resource-acquisition and resource-conservation strategies, consistent with established leaf and root economics frameworks (Wright et al. 2004; Reich 2014; Roumet et al. 2016; Bergmann et al. 2020). Leaf traits included leaf dry matter content, specific leaf area, leaf carbon concentration, and leaf nitrogen concentration. Root traits included root diameter, specific root length, root tissue density, specific root area, root carbon concentration, and root nitrogen concentration. Precipitation treatment was included as an environmental predictor, given the well-documented influence of water availability on AMF functioning and plant responsiveness (Hoeksema et al. 2010; Veresoglou et al. 2012).

### 2.4. Response variable

Plant responsiveness to AMF was quantified as total mycorrhizal growth response, the relative change in plant growth attributable to AMF inoculation. MGR was used in preference to colonization intensity as the primary response variable because the two are frequently decoupled: a plant can be heavily colonized without a correspondingly large growth benefit, or vice versa (Smith and Smith 2011; Hart and Reader 2002). The full Trait2Myco model incorporated ten leaf and root functional traits together with precipitation as predictor variables (Table 1).

**Table 1.**
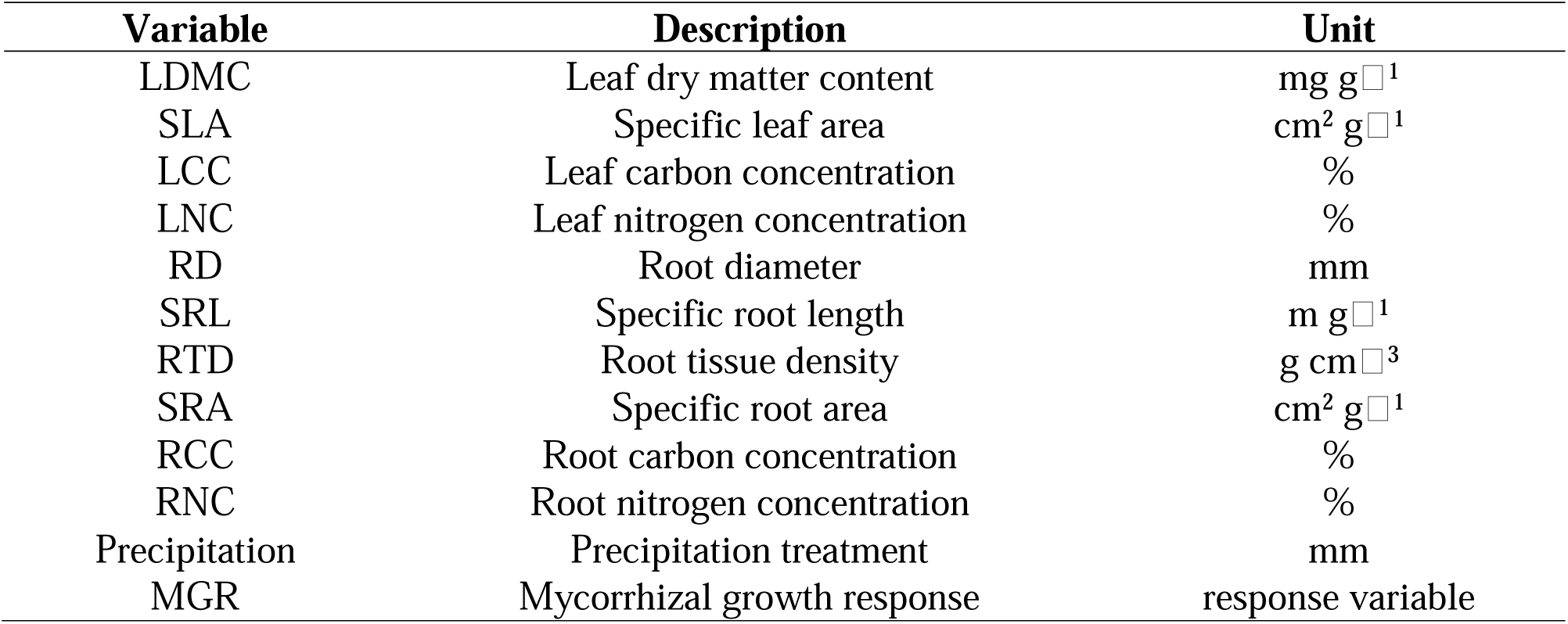
Functional trait and predictor variables incorporated in the Trait2Myco framework.

### 2.5. Random Forest model development

The full Trait2Myco model incorporated all ten functional traits and precipitation as predictors of total MGR. Models were fitted using 1000 trees with variable importance estimation enabled, using the randomForest package in R (Liaw and Wiener 2002).

### 2.6. Variable importance analysis

Predictor importance was quantified as the percentage increase in mean squared error (%IncMSE) under permutation-based variable importance analysis, in which predictors producing larger increases in error following permutation were interpreted as exerting a stronger influence on model performance. Specific leaf area, precipitation, root carbon concentration, and specific root area emerged as the strongest determinants of AMF responsiveness in the full model.

### 2.7. Development of a reduced ecological model

To identify a minimal, biologically interpretable predictor set, a second, reduced model was constructed using only the four variables identified as most influential in the full model: specific leaf area, specific root area, root carbon concentration, and precipitation. This reduced model tested whether a more parsimonious trait set could explain a comparable proportion of variance in MGR relative to the full ten-predictor model.

### 2.8. Training and testing procedure and cross validation

The merged dataset consisted of 80 observations and was randomly partitioned into training (80%; n = 64) and testing (20%; n = 16) subsets using the createDataPartition() function in the caret package (Kuhn 2008). The training dataset was used for model fitting, whereas the testing dataset was held out entirely and used only for evaluating predictive performance on previously unseen observations.

To evaluate whether trait-based prediction generalizes beyond the primary dataset, Trait2Myco was additionally applied to an independent dataset relating host growth traits like plant length, total plant biomass, aboveground dry weight, flower number, and root dry weight to AMF root colonization score (n = 8; Werner and Kiers 2015; https://doi.org/10.5061/dryad.08c2k). A Random Forest model was fitted using the approach described above, with colonization score as the response variable, and evaluated using R², RMSE, and MAE. Variable importance and partial dependence analyses were conducted as for the primary model. Full results of this validation analysis are reported in the Supplementary Material.

Model robustness was additionally assessed using ten-fold cross-validation applied to the full model, implemented through the caret package (Kuhn 2008). Because cross-validation folds were assigned randomly, observations originating from the same species × precipitation combination could occur in different folds. Consequently, cross-validation results represent model consistency across resampled observations but may provide an optimistic estimate of predictive performance owing to biological similarity among replicate observations. Cross-validation of the full model yielded an R² of 0.983, RMSE of 0.046, and MAE of 0.036. Because the full and reduced models were both evaluated on the same independent testing dataset (R² = 0.41 and R² = 0.517, respectively), these held-out test-set values rather than the cross-validation result were considered the more conservative and primary estimates of predictive generalization.

### 2.9. Partial dependence analysis

Partial dependence plots (PDPs) were generated using the pdp package (Greenwell 2017) to estimate the marginal effect of individual predictors on model output while averaging over all remaining predictors. Analyses focused on specific leaf area, root carbon concentration, and precipitation, as these variables consistently ranked among the most important predictors in both the full and reduced models. Principal component analysis (PCA) was performed on the functional trait matrix to visualize multivariate trait space and to assess whether highly responsive and weakly responsive plants occupied distinct regions of that space.

### 2.10. Software availability

Trait2Myco (v0.1.0) is an open-source R package for predicting mycorrhizal growth response (MGR) from plant functional traits and environmental variables. The package is available at https://github.com/tariq6640/Trait2Myco.

## 3. Results

### 3.1. Model performance

Model performance differed among validation approaches, with cross-validation producing higher performance estimates than independent testing (Table 2). The Trait2Myco framework integrated plant functional traits and precipitation data to predict mycorrhizal growth response and identify ecological drivers of responsiveness (Fig. 1). The full Random Forest model, incorporating all ten functional traits and precipitation, explained 41% of variance in mycorrhizal growth response on the independent test set (R² = 0.41, n = 16; Fig. 2), with predicted values showing reasonable agreement with observed MGR across the range of responses, though with greater scatter at higher observed response values. The reduced ecological model, restricted to the four most influential predictors (specific leaf area, specific root area, root carbon concentration, and precipitation), was evaluated on the same held-out test set and achieved improved performance (R² = 0.517), indicating that this smaller, ecologically interpretable predictor set generalized better to unseen observations than the full ten-predictor model. Ten-fold cross-validation of the full model on the complete dataset yielded a substantially higher R² (0.983); because replicate observations within species × precipitation combinations could be distributed across different folds, this value likely overestimates true predictive accuracy and is interpreted as an optimistic upper bound rather than an unbiased performance estimate. The two independent test-set values (R² = 0.41 and R² = 0.517) are therefore treated as the primary, more conservative measures of model generalization throughout. Although ten-fold cross-validation produced very high predictive performance, independent test-set performance was considerably lower, suggesting that cross-validation likely represents an optimistic estimate of predictive accuracy under the structured species × precipitation design (Table 2).

**Figure 2.**
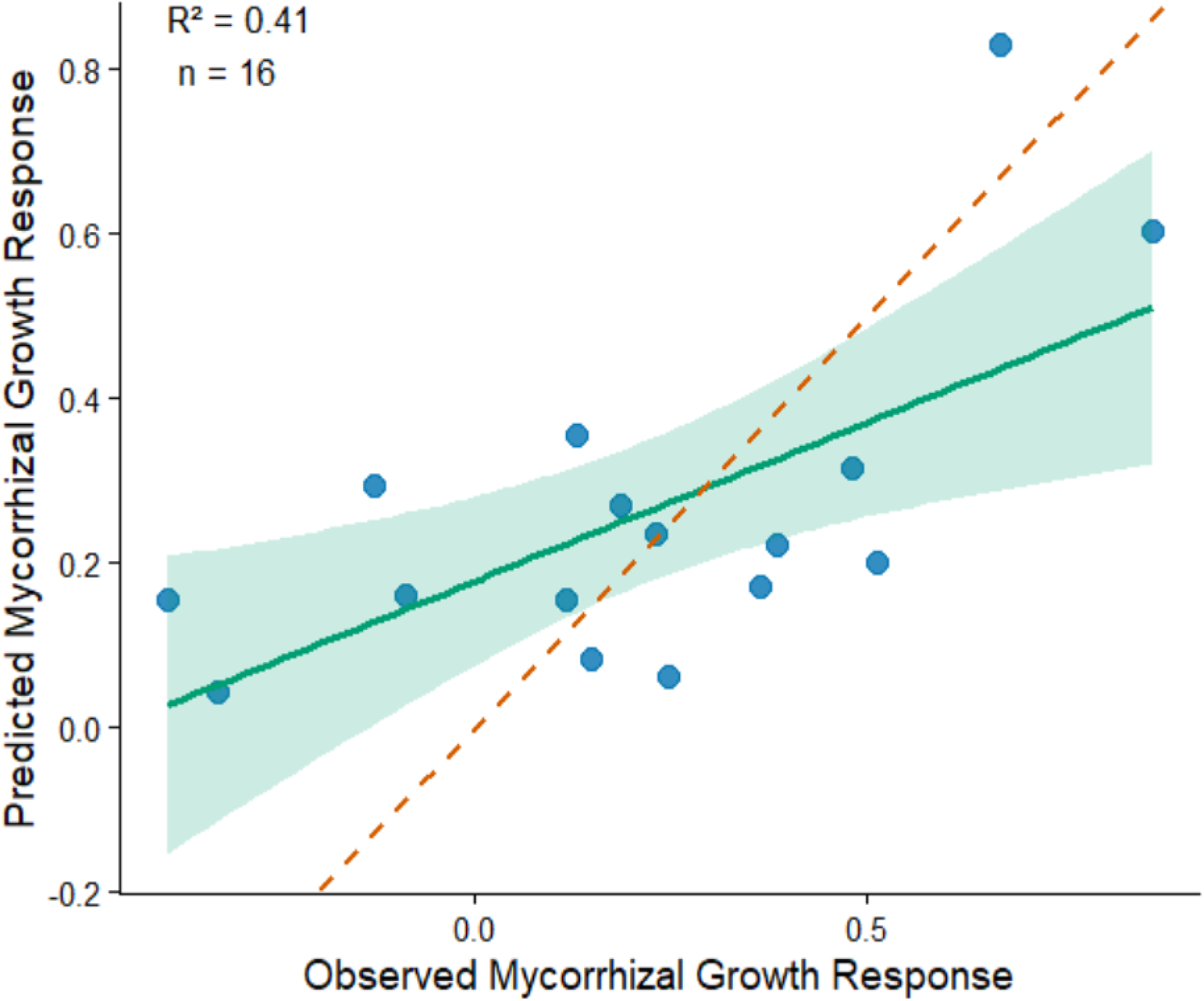
Performance of the Trait2Myco Random Forest model for predicting mycorrhizal growth response (MGR). Predicted values are plotted against observed MGR values for the independent testing dataset (n = 16). The solid green line represents the fitted relationship between observed and predicted values, with the shaded region indicating the 95% confidence interval. The dashed line represents the 1:1 relationship.

**Table 2.** Performance of the Trait2Myco Random Forest models under different validation approaches.

| Model | Predictors | Validation Strategy | R<sup>2</sup> |
| --- | --- | --- | --- |
| Full model | 10 traits + precipitation | Independent test set | 0.41 |
| Reduced model | SLA + SRA + RCC + precipitation | Independent test set | 0.517 |
| Full model | 10 traits + precipitation | 10-fold cross-validation | 0.983 |

SLA: Specific leaf area, SRA: Specific root area, RCC: Root carbon concentration

### 3.2. Variable importance

Permutation-based variable importance analysis identified specific leaf area as the single strongest predictor of mycorrhizal growth response in the full model, followed by precipitation and root carbon concentration (Fig. 3). Specific root area, leaf carbon concentration, leaf nitrogen concentration, root nitrogen concentration, specific root length, root tissue density, and root diameter contributed progressively smaller increases in model error upon permutation, with leaf dry matter content contributing least among the ten traits examined. Both leaf-level and root-level traits appeared among the top predictors, indicating that responsiveness to AMF is not governed solely by either above- or belowground economic strategy.

**Figure 3.**
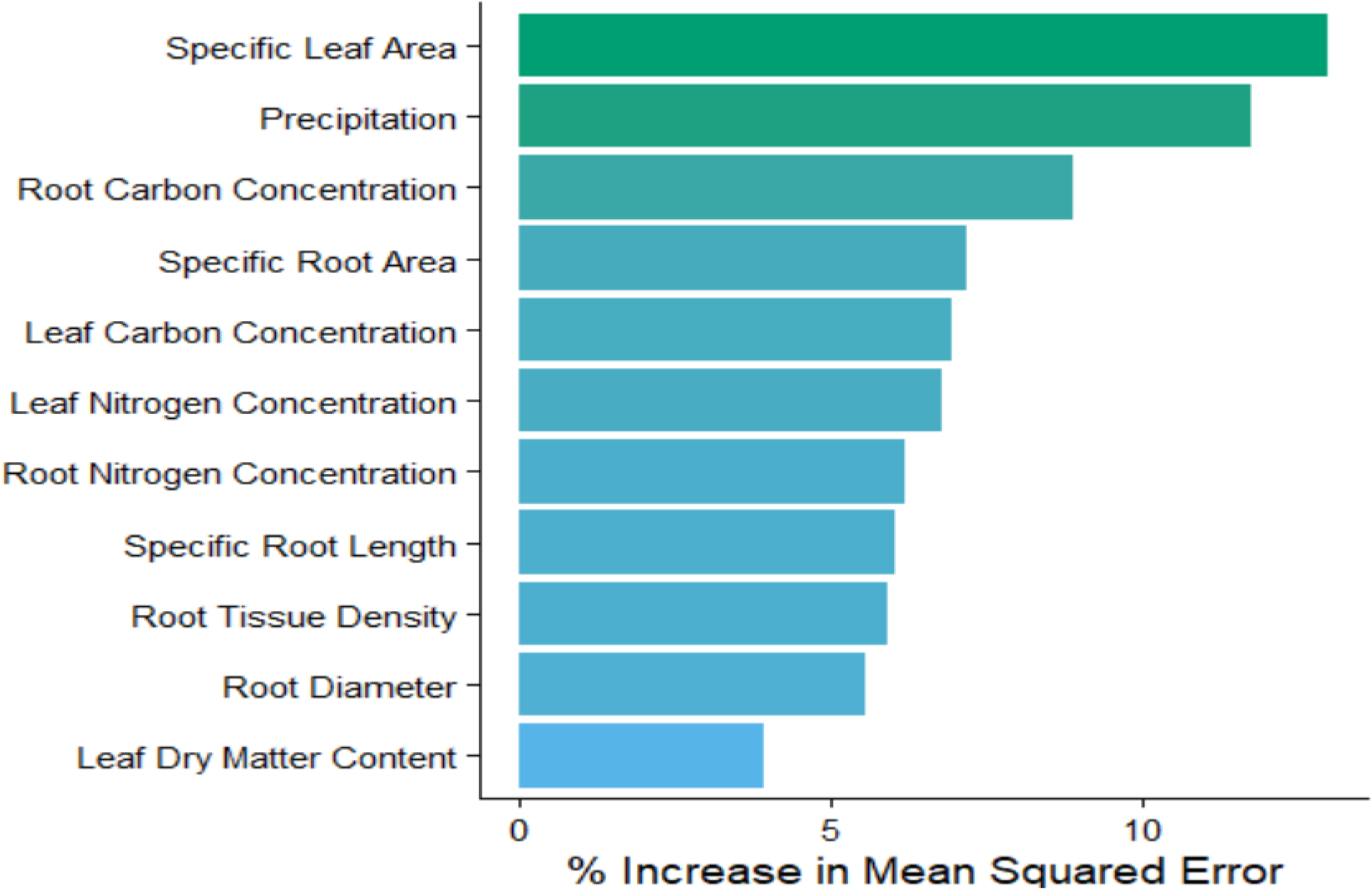
Variable importance ranking from the full Trait2Myco Random Forest model. Predictor importance was quantified as the percentage increase in mean squared error (%IncMSE) following permutation of individual variables. Specific Leaf Area was identified as the strongest predictor of mycorrhizal growth response, followed by precipitation, root carbon concentration, and specific root area. Lower-ranked variables included root diameter, root tissue density, specific root length, and leaf dry matter content. Higher %IncMSE values indicate greater influence on model performance.

### 3.3. Partial dependence on key predictors

Partial dependence plots for the three most important continuous predictors revealed distinct functional relationships with predicted MGR (Fig. 4). Predicted MGR declined sharply with increasing specific leaf area before plateauing at higher specific leaf area values, consistent with reduced mycorrhizal responsiveness among resource-acquisitive, fast-growing species. Predicted MGR showed a unimodal relationship with precipitation, peaking at intermediate rainfall (∼800 mm) and declining at both lower and higher precipitation levels. Root carbon concentration showed a threshold-like relationship with predicted MGR, with a marked increase in response above approximately 4.35% root carbon concentration, beyond which predicted MGR remained comparatively stable across the remaining range of the predictor.

**Figure 4.**
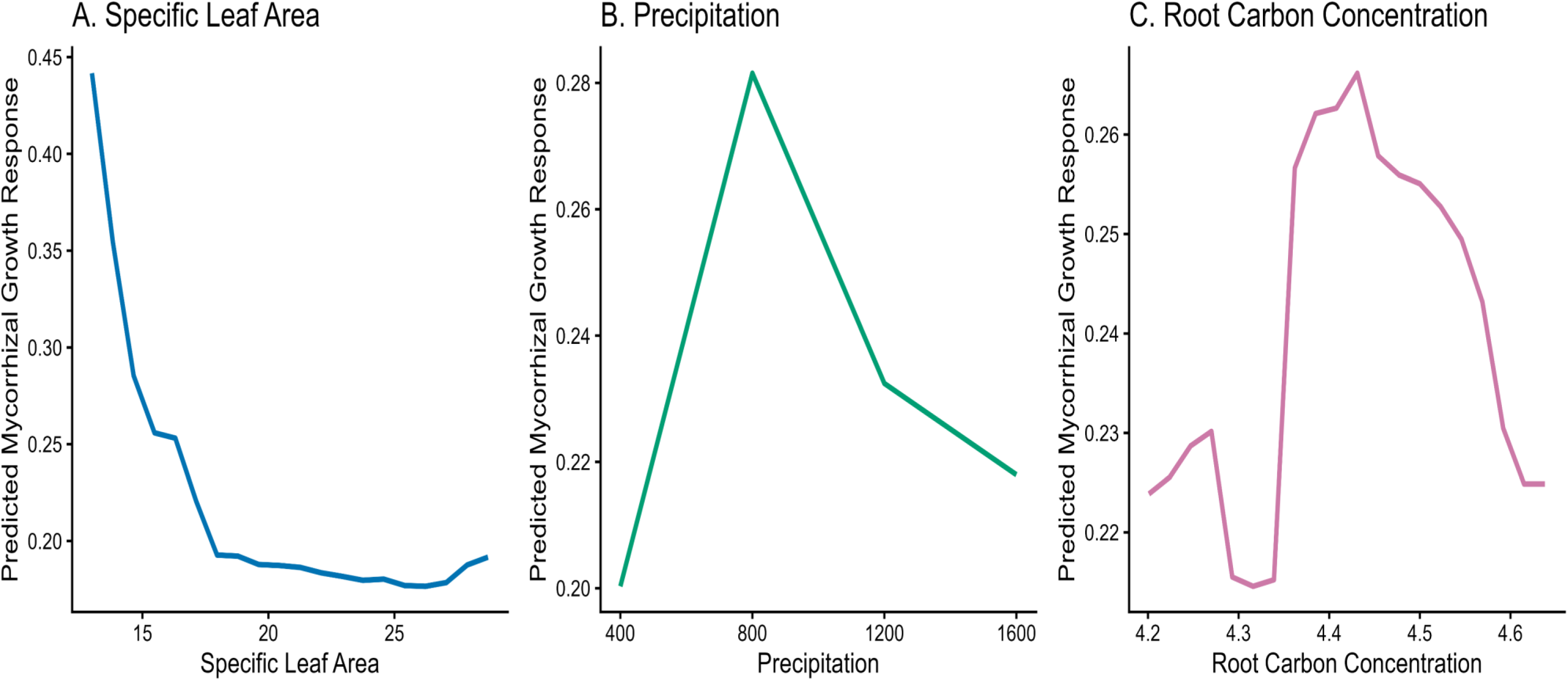
Partial dependence plots illustrating the marginal effects of the three most influential predictors on predicted mycorrhizal growth response (MGR). (A) Specific Leaf Area exhibited a strong negative relationship with MGR, with responsiveness declining rapidly across the observed specific leaf area gradient before reaching a plateau. (B) Precipitation showed a unimodal relationship with MGR. (C) Root Carbon Concentration displayed a threshold-like response, with predicted MGR increasing sharply above approximately 4.35% root carbon concentration and remaining relatively stable thereafter. Partial dependence plots depict the effect of each predictor while averaging over the effects of all remaining variables in the model.

### 3.4. Relative contribution of trait categories

Aggregating variable importance scores by category showed that leaf traits and root traits contributed almost equally to overall model performance, accounting for 41.9% and 41.3% of cumulative variable importance, respectively, while precipitation (environmental factors) accounted for the remaining 16.8% (Fig. 6). This near-parity between leaf and root trait contributions indicates that above- and belowground economic strategies jointly, rather than independently, structure variation in mycorrhizal growth response in this dataset.

### 3.5. Multivariate trait space

Principal component analysis of the functional trait matrix showed that PC1 and PC2 together captured the dominant axes of trait variation (Fig. 5). Individuals with strongly positive mycorrhizal growth response were concentrated toward the negative end of PC1, in the direction associated with root carbon concentration, while individuals with weaker or negative response were more broadly distributed toward the positive end of PC1, in the direction associated with specific root length, specific root area, specific leaf area, and leaf nitrogen concentration. This pattern indicates partial separation of highly responsive and weakly responsive observations along a multivariate trait axis consistent with variation in plant resource-acquisition and resource-conservation strategies.

**Figure 5.**
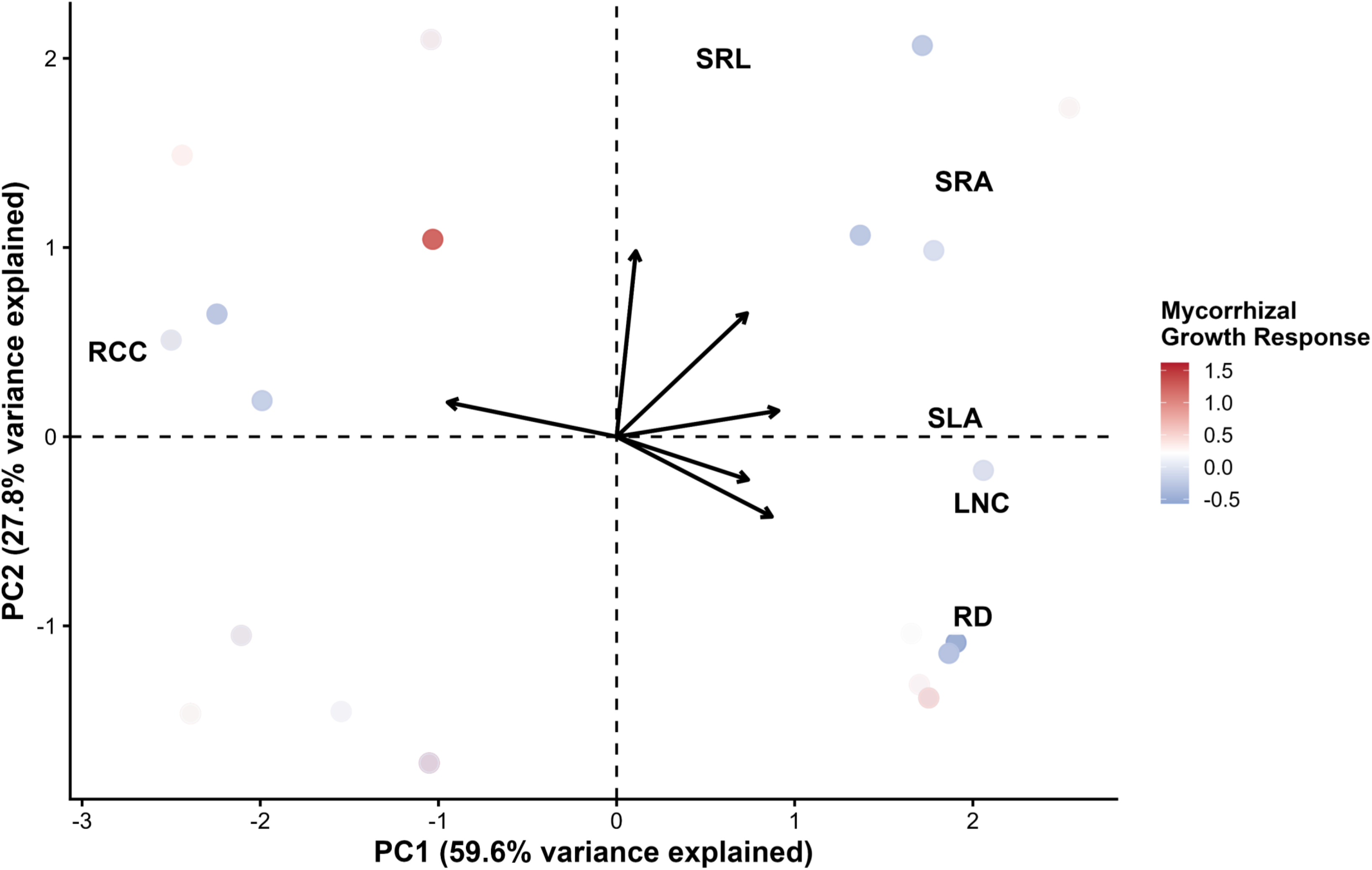
Principal component analysis (PCA) of plant functional traits associated with responsiveness to arbuscular mycorrhizal fungi. Points represent individual observations and are coloured according to total mycorrhizal growth response (MGR), with warmer colours indicating greater responsiveness. Trait vectors represent the direction and magnitude of trait contributions to multivariate variation.

**Figure 6.**
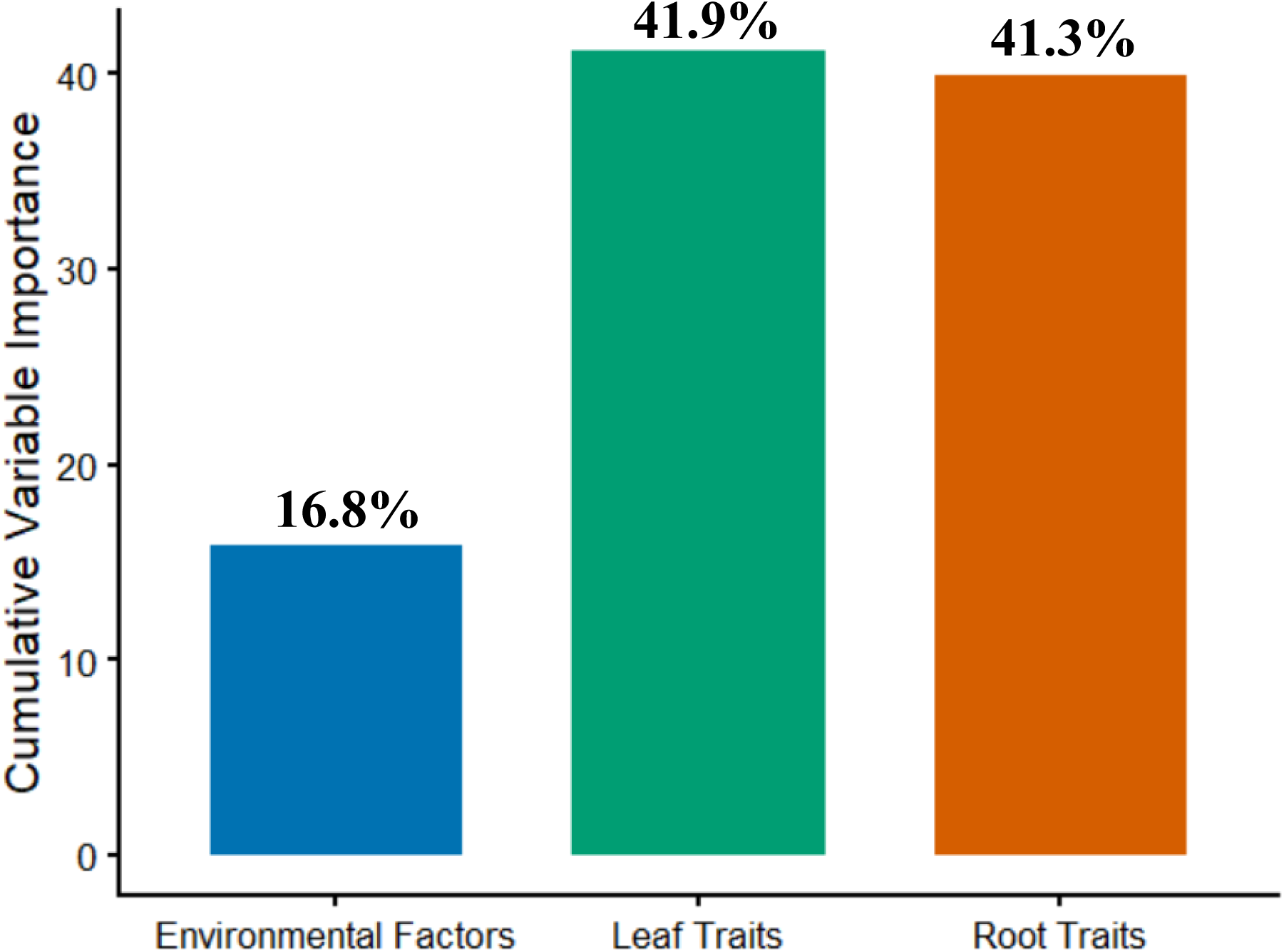
Relative contribution of predictor categories to model performance in the Trait2Myco framework. Variable importance values were aggregated into three ecological categories: environmental factors (precipitation), leaf traits, and root traits. Leaf traits contributed 41.9% of cumulative model importance, root traits contributed 41.3%, and environmental factors contributed 16.8%

### 3.6. Supplementary response trait correlation

Pairwise correlation analysis of all functional traits against total MGR (Fig. S1) showed that specific leaf area had the strongest negative correlation with MGR among all traits examined, followed by leaf nitrogen concentration and root diameter, while root carbon concentration and root tissue density showed the strongest positive correlations with MGR (Fig. S2). These univariate patterns were broadly consistent with the variable importance rankings and partial dependence relationships identified by the Random Forest model, providing convergent support for specific leaf area and root carbon-related traits as the dominant correlates of mycorrhizal responsiveness.

### 3.7. External validation: predicating AMF colonization from plant growth traits

To evaluate whether a Random Forest, trait-based approach generalizes beyond MGR prediction, a second model was trained on an independent dataset relating plant growth traits to AMF root colonization score (Werner and Kiers 2015). This model achieved strong predictive performance on a small independent sample (R² = 0.94, RMSE = 0.070, MAE = 0.054, n = 8; Fig. S3), with predicted colonization scores closely tracking observed values across the full range of colonization intensity. Plant length was the strongest predictor of colonization score, followed closely by total plant biomass and aboveground dry weight, with flower number and root dry weight contributing comparatively less (Fig. S3). Partial dependence plots showed that predicted colonization increased sharply with plant length up to approximately 10 cm before plateauing and showed a similar threshold-like increase with total plant biomass and root dry weight above intermediate values (Fig. S4). Pairwise correlations among growth traits and colonization score were uniformly positive and strong (r = 0.62–0.85; Fig. S5), consistent with the pattern that more vigorously growing plants supported higher colonization intensity in this dataset. Given the small sample size (n = 8), these results should be interpreted as a preliminary demonstration of cross-dataset generalizability rather than a definitive validation and are reported in full in the Supplementary Material.

## 4. Discussion

Although AMF colonization is frequently used as a proxy for symbiotic success, colonization intensity and host benefit are often decoupled. Consequently, understanding the determinants of mycorrhizal growth remains a central challenge in mycorrhizal ecology. While AMF associations are widespread across terrestrial ecosystems, the magnitude of plant response varies substantially among species and environments. In this study, we developed Trait2Myco, a Random Forest–based framework that integrates plant functional traits and environmental variables to predict MGR. The framework identified specific leaf area, precipitation, root carbon concentration, and specific root area as the strongest predictors of responsiveness, suggesting that AMF benefits are linked to coordinated plant economic strategies rather than isolated trait effects.

### 4.1. Resource-acquisitive trait syndromes influence responsiveness to AMF

Specific leaf area emerged as the strongest predictor of MGR in both the full and reduced Trait2Myco models. Specific leaf area is a central component of the plant economics spectrum and is widely associated with rapid resource acquisition, high photosynthetic capacity, and greater nutrient demand (Wright et al. 2004; Reich 2014). Species characterized by high specific leaf area generally acquire resources quickly and maintain fast growth rates, whereas species with lower specific leaf area invest in denser and longer-lived tissues. The negative relationship between specific leaf area and predicted responsiveness observed in this study suggests that more acquisitive species may derive relatively lower benefits from AMF within the ecological space represented by our dataset.

This pattern is broadly consistent with ecological theories proposing that plants with efficient direct nutrient acquisition pathways may rely less heavily on fungal nutrient delivery (Koide 1991; Chagnon et al. 2013). In contrast, species exhibiting more conservative strategies may obtain greater returns from fungal-mediated nutrient acquisition because symbiotic pathways complement slower intrinsic resource-acquisition strategies. Importantly, because functional traits and responsiveness were measured in separate but linked experimental datasets, these findings should be interpreted as species-level associations rather than direct causal relationships. Nevertheless, the recovery of specific leaf area as the most influential predictor by Trait2Myco suggests that leaf economic strategies are strongly associated with variation in AMF responsiveness.

### 4.2. Root carbon concentration as a previously underappreciated predictor

One of the most interesting findings of this study was the strong influence of root carbon concentration. While traits such as root diameter, specific root length, and specific root area are commonly discussed in studies of plant AMF interactions, root carbon concentration has rarely been evaluated explicitly as a predictor of responsiveness. Because carbon represents the principal resource supplied by plants to fungal partners, variation in belowground carbon investment may directly influence the costs and potential benefits of maintaining symbiosis (Smith and Read 2008; Kiers et al. 2011; Fellbaum et al. 2012). Experimental evidence indicates that carbon supplied by the host can directly regulate fungal nutrient uptake and transport processes, suggesting a mechanistic link between host carbon economy and symbiotic benefit (Fellbaum et al. 2012). The strong influence of root carbon concentration in Trait2Myco therefore supports the hypothesis that carbon allocation represents a key determinant of AMF responsiveness.

The threshold-like increase in predicted MGR at higher root carbon concentration values suggests that carbon economy may play a more important role in determining responsiveness than previously recognized. Plants allocating larger carbon pools belowground may be better able to support fungal metabolism and sustain nutrient-exchange structures, thereby increasing the net benefits derived from AMF associations. Although mechanistic validation will be required, the identification of root carbon concentration as a major predictor highlights a potentially overlooked component of mycorrhizal ecology and suggests promising avenues for future research linking carbon allocation strategies with symbiotic performance.

### 4.3. Root traits contribute substantially to mycorrhizal outcomes

Specific root area was consistently identified among the most important predictors of responsiveness. Root morphological traits influence nutrient foraging efficiency, soil exploration, and opportunities for fungal colonization (Comas et al. 2013; Roumet et al. 2016). Species exhibiting higher specific root area and specific root length generally possess greater soil exploration capacity and resource acquisition potential, traits frequently associated with variation in belowground interactions (Comas et al. 2013). Plants exhibiting greater root surface area may simultaneously increase direct nutrient uptake and enhance access to AMF-derived nutrient pathways. The importance of specific root area suggests that root economic strategies contribute significantly to determining whether plants benefit from mycorrhizal partnerships.

Interestingly, aggregated importance scores demonstrated that leaf traits (41.9%) and root traits (41.3%) contributed almost equally to overall model performance. This finding indicates that responsiveness reflects whole-plant strategies rather than exclusively aboveground or belowground processes. Recent studies have emphasized that root and leaf economics are only partially coordinated and should be considered together when evaluating plant responses to environmental change and biotic interactions (Bergmann et al. 2020; Freschet et al. 2021). Our results support this perspective and suggest that AMF responsiveness emerges from integrated plant economic syndromes.

### 4.4. Environmental context strongly modifies responsiveness

Precipitation was the second most influential predictor overall and the most important environmental factor in the model. Water availability affects AMF activity, nutrient mobility, root growth, and plant carbon allocation, all of which influence the costs and benefits of symbiosis (Smith and Read 2008; van der Heijden et al. 2015). Partial dependence analyses demonstrated a nonlinear relationship between precipitation and MGR, with responsiveness peaking at intermediate precipitation levels and declining at both wetter and drier extremes.

This pattern is consistent with the view that AMF outcomes are strongly context dependent (Johnson et al. 1997; Hoeksema et al. 2010). Under high water availability, plants may acquire resources effectively without fungal assistance, reducing the relative value of symbiosis. Conversely, under severe drought, host carbon limitation and reduced fungal activity may constrain the benefits of AMF associations. Maximum responsiveness is therefore expected under intermediate environmental conditions where resource limitation creates opportunities for fungal contributions while plant carbon supply remains sufficient to sustain the symbiosis.

### 4.5. Functional trait syndromes explain responsiveness better than individual traits

The PCA revealed that highly responsive and weakly responsive observations occupied different regions of multivariate trait space. Observations associated with greater MGR were positioned along gradients defined by specific leaf area, specific root area, specific root length, root carbon concentration, root diameter, and leaf nitrogen content, indicating that responsiveness is associated with combinations of traits rather than single predictors. Recent work has emphasized that root trait variation is multidimensional and cannot be adequately represented by a single axis of resource acquisition, highlighting the importance of trait syndromes rather than isolated traits (Weemstra et al. 2016; Bergmann et al. 2020). The clustering patterns observed in the PCA therefore support the idea that mycorrhizal responsiveness emerges from coordinated suites of leaf and root traits. This pattern reinforces the notion that plant AMF interactions should be interpreted within the broader context of coordinated resource-acquisition strategies.

The PCA findings closely complement the Random Forest analyses. Whereas Random Forest identified the most influential predictors, PCA demonstrated how these traits collectively structure ecological variation among observations. Together, these analyses suggest that AMF responsiveness is an emergent property of integrated leaf and root trait syndromes rather than the consequence of individual trait effects acting independently.

### 4.6. Trait2Myco as a predictive ecological framework

An important contribution of this study is the development of Trait2Myco as a predictive framework rather than merely a statistical model. Trait2Myco successfully identified functional trait syndromes associated with AMF responsiveness while also detecting nonlinear relationships that may be difficult to recover using traditional statistical approaches. The reduced ecological model retained substantial predictive power despite using only four predictors, demonstrating that meaningful prediction is possible using relatively small trait sets.

Importantly, Trait2Myco should be viewed as a framework rather than a novel machine-learning algorithm. The novelty lies in its ecological application, integration of functional trait theory with predictive modeling, and ability to identify trait syndromes associated with responsiveness. Because the framework uses readily measurable variables, it may provide practical utility for plant breeding, microbial inoculant development, restoration ecology, and agricultural decision-making.

## 5. Limitations

First, trait measurements and mycorrhizal response measurements originated from separate but linked experimental datasets. Consequently, trait values were effectively constant across precipitation treatments within species, and the effective number of unique trait syndromes remained relatively small. This means the model should be interpreted primarily as identifying species-level trait associations rather than individual-level causal predictors.

Second, only four plant species were included in the dataset. Although the framework identified biologically coherent relationships, broader taxonomic representation will be necessary to evaluate the generality of these patterns. Third, the substantial difference between cross-validation performance (R² = 0.983) and independent test-set performance (R² = 0.517) likely reflects biological similarities among replicate observations. While cross-validation demonstrates strong predictive signal, the test-set results should be considered the more conservative estimate of model transferability.

Finally, Trait2Myco focused on plant traits and precipitation and did not incorporate fungal identity, soil properties, microbial community composition, or nutrient availability. Inclusion of additional biotic and abiotic variables may improve predictive performance and broaden ecological applicability.

## 6. Conclusions

Trait2Myco demonstrates that plant responsiveness to arbuscular mycorrhizal fungi can be predicted using a relatively small number of functional traits and environmental variables. Specific leaf area, root carbon concentration, specific root area, and precipitation emerged as dominant predictors of responsiveness, indicating that both leaf and root economic strategies contribute to determining AMF outcomes. By integrating machine learning with ecological theory, Trait2Myco provides a reproducible framework for identifying functional trait syndromes associated with mycorrhizal responsiveness and offers a foundation for future efforts aimed at predicting plant–fungus interactions across environmental gradients.

## Data availability statement

The Trait2Myco source code, trained Random Forest model, package documentation, and example workflows are publicly available through the GitHub repository Trait2Myco at https://github.com/tariq6640/Trait2Myco. The primary datasets used in this study were obtained from publicly available Dryad repositories and are cited in the manuscript. Trait2Myco can be installed directly in R from GitHub using the remotes::install_github() function.

