## Supplemental materials for "Machine Learning Identification of Functional Trait Syndromes Associated with Responsiveness to Arbuscular Mycorrhizal Fungi"

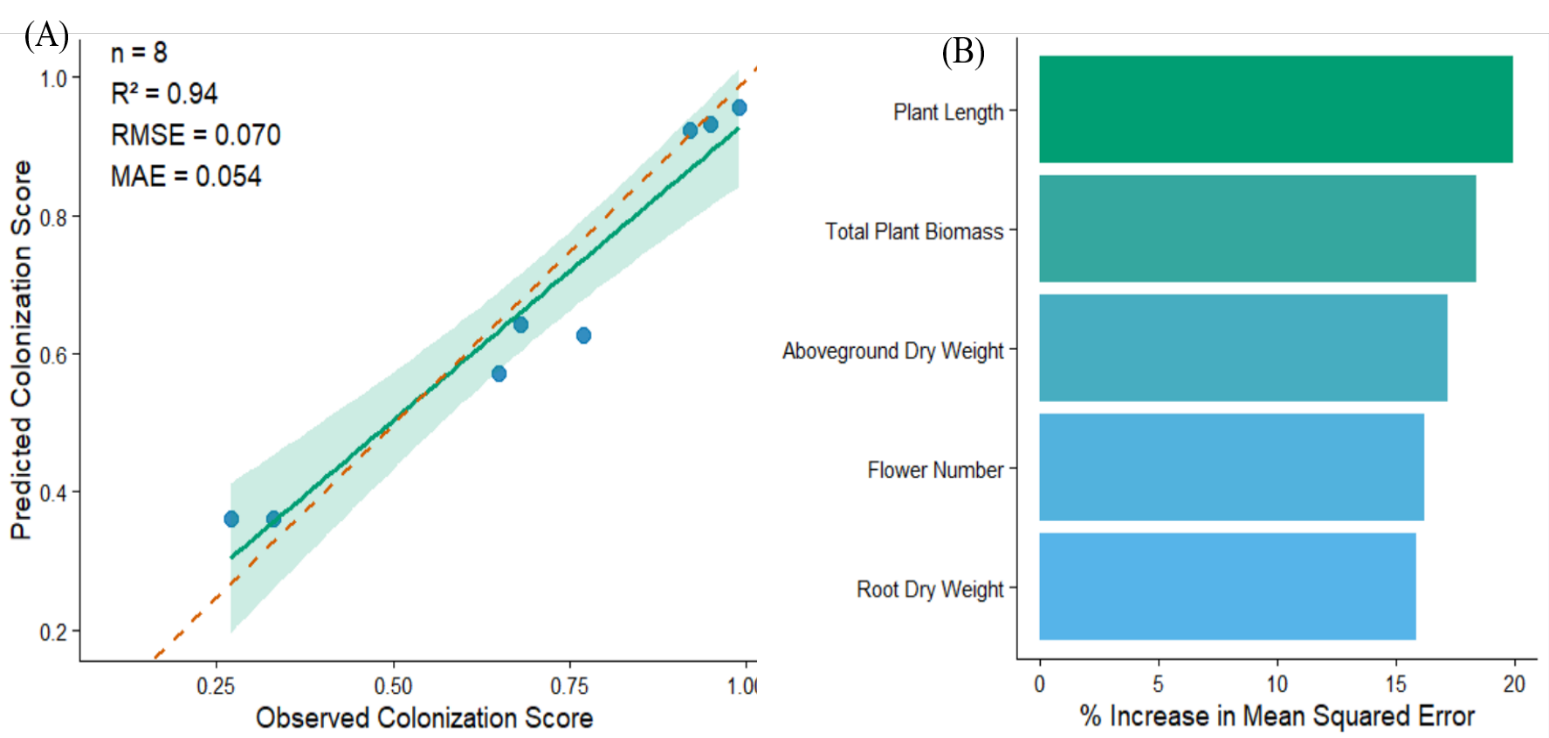

**Figure S1.** Observed versus predicted AMF colonization scores from the supplementary Random Forest validation model based on plant growth traits.

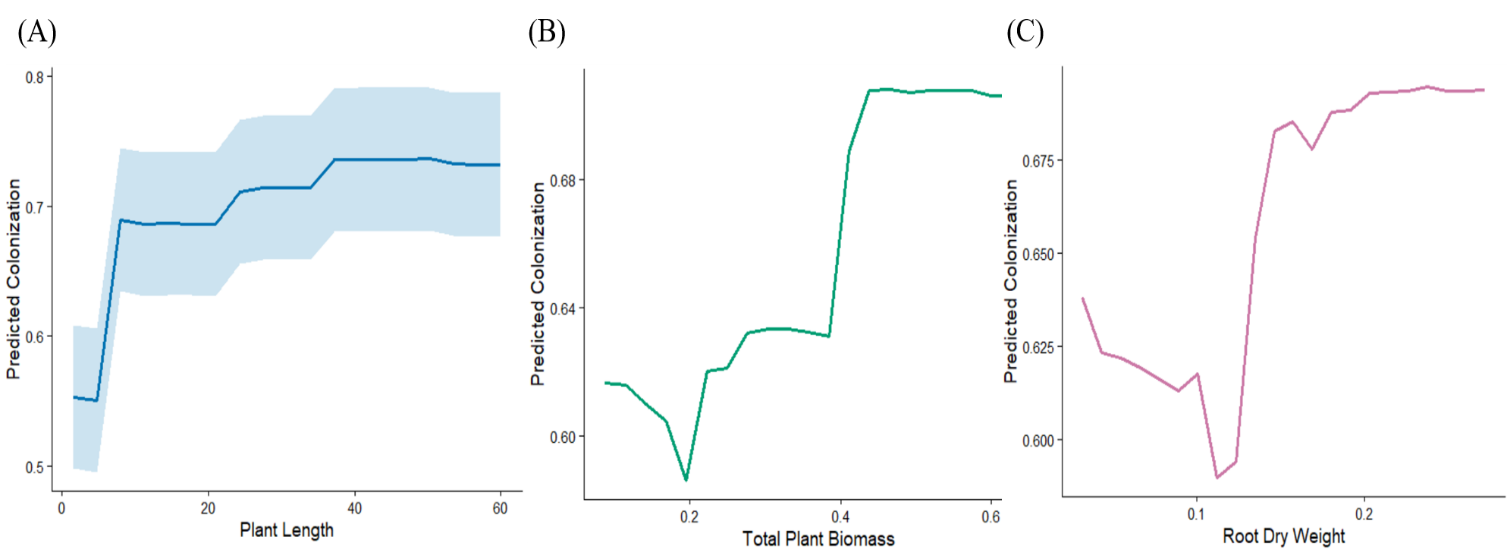

**Figure S2.** Variable importance ranking for predictors of AMF colonization in the supplementary validation model.

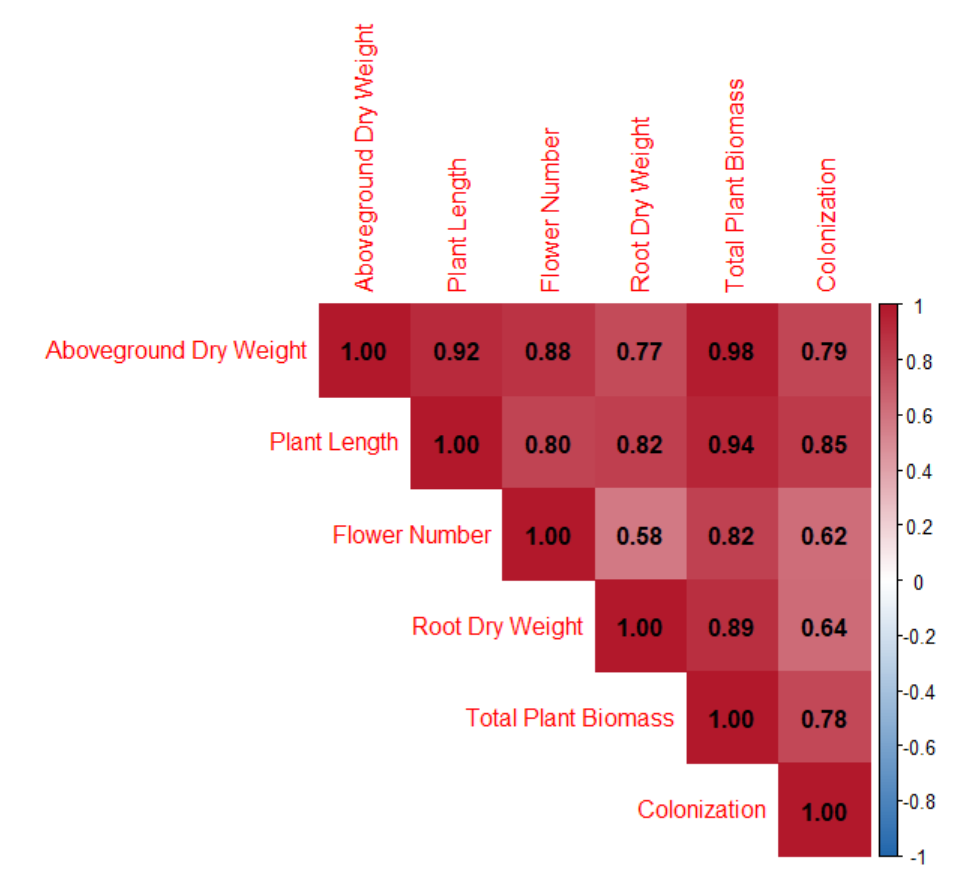

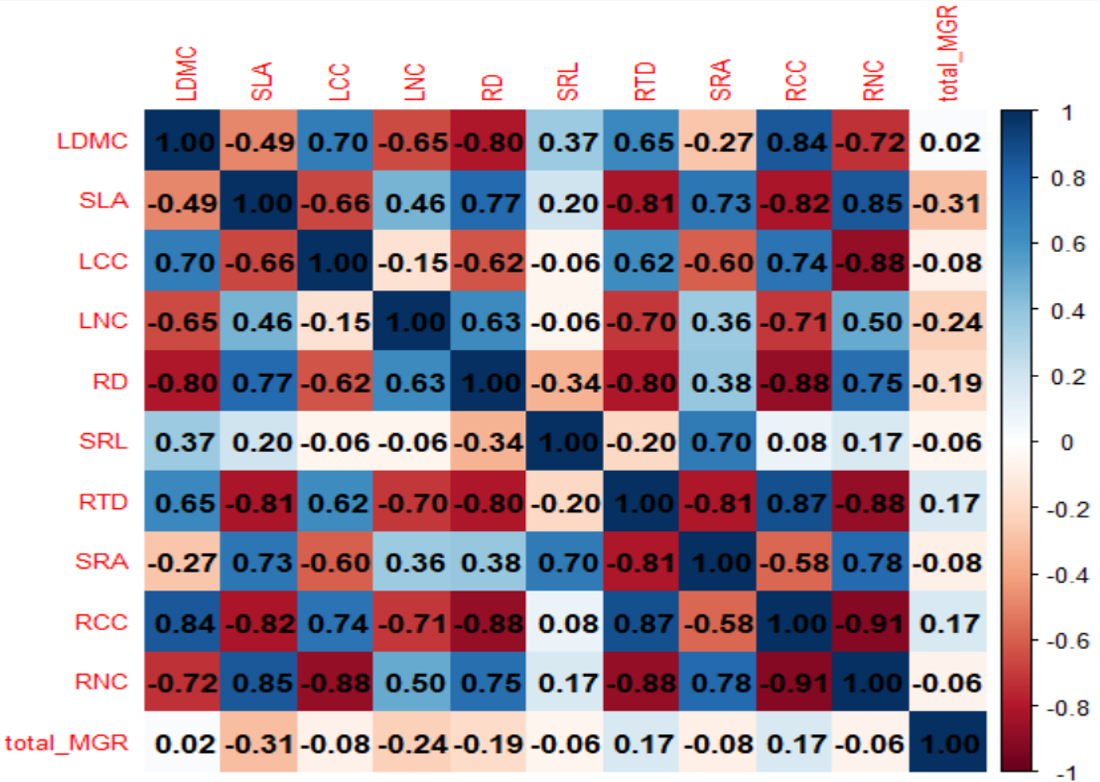
**Figure S3.** Partial dependence plots illustrating the effects of plant length, total biomass, and root biomass on predicted AMF colonization.

**Figure S4.** Correlation matrix of plant functional traits and mycorrhizal growth response used in the Trait2Myco framework.

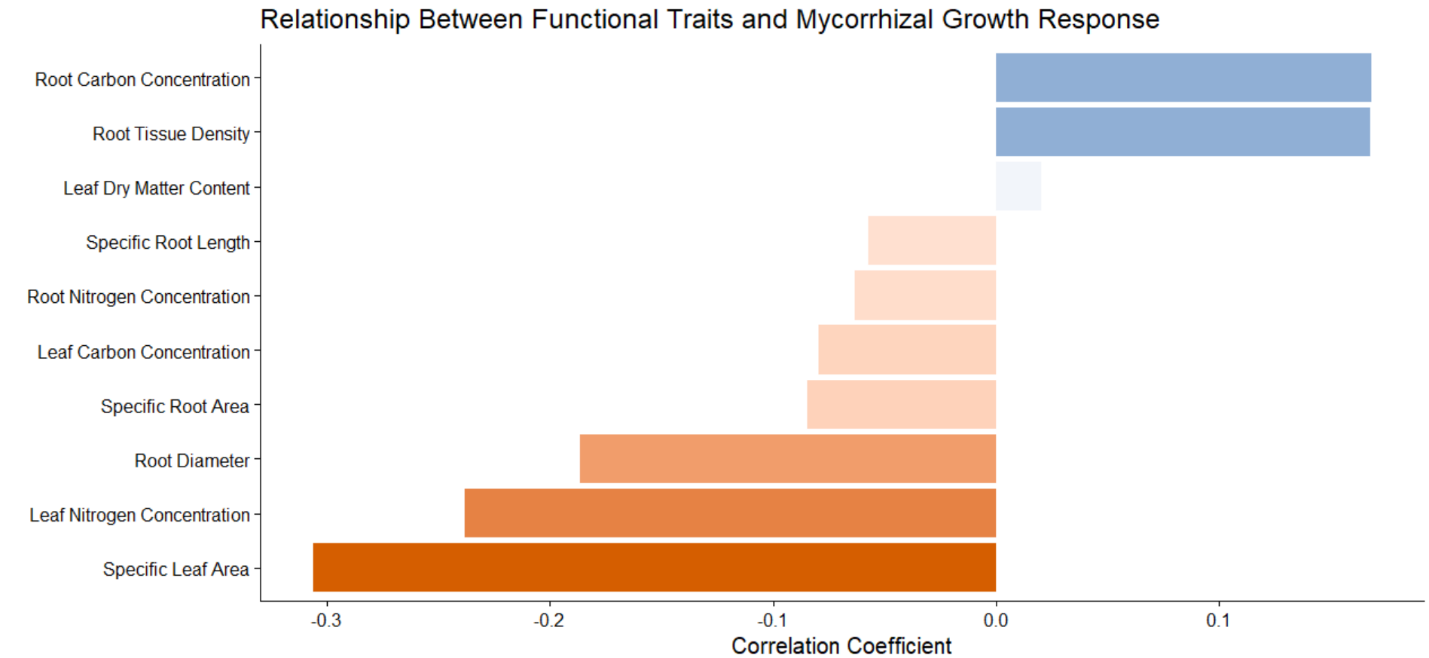

**Figure S5.** Relationships among plant growth traits and AMF colonization intensity in the independent validation dataset.
